# A Rab Escort Protein Regulates the MAPK Pathway That Controls Filamentous Growth in Yeast

**DOI:** 10.1101/2020.06.02.130690

**Authors:** Sheida Jamalzadeh, Paul J. Cullen

**Affiliations:** Department of Chemical and Biological Engineering, University at Buffalo, State University of New York, Buffalo New York; Department of Biological Sciences, University at Buffalo, State University of New York, Buffalo New York

**Keywords:** Rab Escort Protein, MAP kinase, Cdc42, Protein Trafficking, Cell Polarity, Genomics

## Abstract

MAPK pathways regulate different responses yet can share a subset of common components. In this study, a genome-wide screen was performed to identify genes that, when overexpressed, induce a growth reporter (*FUS1-HIS3*) that responds to ERK-type MAPK pathways (Mating/filamentous growth or fMAPK) but not p38-type MAPK pathways (HOG) in yeast. Approximately 4,500 plasmids overexpressing individual yeast genes were introduced into strains containing the *FUS1-HIS3* reporter by high-throughput transformation. Candidate genes were identified by measuring the degree of growth, which was a reflection of reporter activity. Of fourteen genes identified and validated by re-testing, two were metabolic controls (*HIS3* and *ATR1*), five had established roles in regulating ERK-type pathways (*STE4, STE7, BMH1, BMH2, MIG2*) and seven represent potentially new regulators of MAPK signaling (*RRN6, CIN5, MRS6, KAR2, TFA1, RSC3, RGT2*). *MRS6* encodes a Rab escort protein and effector of the TOR pathway that plays an established role in nutrient signaling. *MRS6* overexpression stimulated filamentous/invasive growth and phosphorylation of the ERK-type fMAPK, Kss1. Overexpression of *MRS6* reduced the osmotolerance of cells and phosphorylation of the p38/HOG pathway MAPK, Hog1. Mrs6 interacted with the PAK kinase Ste20 and MAPKK Ste7 by two-hybrid analysis. Collectively, the data indicate that Mrs6 may function to selectively propagate an ERK-dependent signal. Generally speaking, the identification of new MAPK pathway regulators by genetic screening in yeast may be a useful resource for understanding signaling pathway regulation.

## INTRODUCTION

During cell differentiation, cells specialize into specific types by the action of signal transduction pathways. <u>M</u>itogen-<u>a</u>ctivated <u>p</u>rotein <u>k</u>inase (MAPK) pathways control numerous responses, including cell differentiation, proliferation, cell migration, and apoptosis (Sun et al. 2015; Cicenas et al. 2017). MAPK pathways control diverse responses by regulating the expression of a large number of target genes. There are four types of MAPK pathways: RAF-MEK-ERK1/2, JNK1/2/3, p38α/β/γ/δ, and ERK5 (Cicenas et al. 2017; Peng et al. 2018). Remarkably, these pathways can share common components, which leads to proper cross-talk in normal settings and unregulated cross-talk in the disease state. Mis-regulation of MAPK signaling leads to inappropriate responses, such as cancers and problems with immune system function (Silva 2004; Raman et al. 2007; Papa et al. 2019). Due to the crucial roles of MAPK pathways in regulating fundamental cellular processes, they remain the focus of the investigation by many labs and are a focus for therapeutic targeting (Roberts et al. 1997; Alory and Balch 2001; de Dios et al. 2010; Kim and Choi 2015; McGivern et al. 2018; Smalley and Smalley 2018).

MAPK pathways are evolutionarily conserved signalling modules in eukaryotes, and fundamental insights into MAPK pathway regulation have come from studies in many systems. The budding yeast *Saccharomyces cerevisiae* is a unicellular organism that has been extensively used as a model for studying signaling pathways (McCaffrey et al. 1987b; Roberts et al. 1997; Brizzio et al. 1999; Schwartz and Madhani 2004; Dohlman and Slessareva 2006; Karunanithi et al. 2012; Lasserre et al. 2015; Mizuno et al. 2015). Like in other eukaryotes, yeast utilizes ERK-type and p38-type MAPK pathways (Saito 2010b; Martin 2019). One ERK-type pathway mediates the response to nutrient-limiting conditions that permit filamentous (pseudohyphal/invasive) growth, a fungal-type foraging response resulting in the formation of chains of elongated interconnected cells (Gimeno et al. 1992; Roberts and Fink 1994a). This pathway functions through a set of kinases that function in a tandem series: p21 activated [PAK] Ste20 (MAPKKKK), Ste11 (MAPKKK), Ste7 (MAPKK), and Kss1 (MAPK) (Roberts and Fink 1994b; Cullen and Sprague 2000). A second ERK-type pathway in yeast controls the mating of haploid cells through an almost identical set of kinases: Ste20 (PAK), Ste11 (MAPKKK), Ste7 (MAPKK), and Fus3 and Kss1 (MAPK). Two MAPKs, Fus3, and Kss1, function in mating and filamentous growth pathways, respectively. It has been shown that the deletion of *KSS1* causes a reduction in agar penetration (Cook et al. 1997), a phenotype called invasive growth that is related to filamentous growth (Roberts and Fink 1994a), while it has little effect on mating efficiency (Madhani et al. 1997). In contrast, deletion of *FUS3* allows cells to penetrate the agar more vigorously (Cook et al. 1997) while they cause a moderate decrease in mating efficiency (Madhani et al. 1997). This and other data support the idea that one MAPK promotes invasive/filamentous growth (Kss1), and while another mainly functions to regulating mating (Fus3). Surprisingly, the elimination of both MAPKs results in more agar penetration, which identified an inhibitory role for the unphosphorylated form of Kss1 regulating filamentous growth (Cook et al. 1997; Madhani et al. 1997; Bardwell et al. 1998a; Bardwell et al. 1998b).

A p38-type pathway, the high osmolarity glycerol response (HOG) pathway, allows the response to hyperosmotic conditions through Pbs2 (MAPKK) and Hog1 (MAPK) (Brewster et al. 1993; Albertyn et al. 1994; Hohmann 2002). One branch of this pathway shares components with the mating and fMAPK pathways (Posas and Saito 1997). Specifically, Ste20 and Ste11 function to regulate Pbs2 and Hog1. Therefore, MAPK pathways in yeast can share some common components despite the fact that the pathways induce different transcriptional and morphogenetic responses.

In pathogens, the filamentation response is critical for host-cell attachment, invasion into tissues, and virulence (Berman 2006). In *S. cerevisiae* haploid cells, filamentous growth is triggered by growth in a non-preferred carbon source. The response is regulated by multiple signal transduction pathways (Bharucha et al. 2008; Jin et al. 2008), including the RAS-cAMP-PKA pathway (Gimeno et al. 1992; Mosch et al. 1996; Harashima and Heitman 2002; Pan and Heitman 2002) and the filamentous growth MAPK pathway (fMAPK) (Roberts and Fink 1994b). These pathways induce target genes that reorganize cell polarity, the cell cycle, and cell adhesion to bring about a new cell type (Kron 1997; Madhani and Fink 1997; Madhani 2000; Pan et al. 2000). The signaling mucin Msb2 operates at the head of the fMAPK pathway, and through the adaptor protein, Sho1, regulates MAPK activity by interaction with the Ras-homology (Rho)-type GTPase Cdc42. Sho1 interacts with Msb2 and Ste11 and functions in both the fMAPK and HOG pathways (Maeda et al. 1995a; Posas and Saito 1997; Tatebayashi et al. 2006). Cdc42 is an essential gene that is required for the maintenance of cell polarity and signaling. Human homolog Cdc42 is 81% identical to the yeast protein (Adams et al. 1990; Johnson and Pringle 1990; Ziman et al. 1993; Johnson 1999; Pruyne and Bretscher 2000; Kachroo et al. 2015). Cdc42 regulates the fMAPK pathway by interacting with Ste20 (Roberts and Fink 1994b; Cook et al. 1997; Madhani and Fink 1997).

Several mechanisms that promote insulation have been described. One mechanism involves scaffolds, such as Ste5 (Chol et al. 1994; Marcus et al. 1994; Printen and Sprague 1994) and Pbs2 (Maeda et al. 1995b). Ste5 activates Fus3 by forming a multi-kinase complex that joins the Ste11, Ste7, and Fus3 kinases (Chol et al. 1994; Zarrinpar et al. 2004; Bhattacharyya et al. 2006). Pbs2 regulates the HOG pathway by being activated through two different branches, *SLN1-SSK1* and Sho1 (Maeda et al. 1995b). Another mechanism that is employed to maintain specificity involves cross-pathway inhibition. In this case, a transcription factor for the filamentation pathway, Tec1, is phosphorylated by Fus3, which leads to its turnover by a ubiquitin ligase complex (Bao et al. 2004; Chou et al. 2004). An intriguing challenge, therefore, is to understand how pathways that share elements establish and maintain their identity (Good et al. 2011; Witzel et al. 2012).

The core MAPK regulators (MAPKKK->MAPKK->MAPK) are well known, and several proteins have been identified that regulate the fMAPK pathway at or above the level of Cdc42. However, some proteins that regulate these pathways may remain unidentified. For example, genetic and genome-wide screens continue to identify new proteins that regulate the fMAPK pathway (Chavel et al. 2014). We developed a screen to identify new regulators of ERK-type pathways in yeast. By focusing on one new pathway regulator, Mrs6, we identify a possible adaptor-based mechanism that may promote pathway insulation. Since many of the new regulators identified have homologs in other eukaryotes, including humans, investigation of fMAPK pathway regulators provides a foundation for understanding MAPK pathway regulation in general. This may contribute to the development of new therapeutic targets in related species of fungal pathogens and can be linked to other signaling systems in higher organisms, with implications for human disease.

## RESULTS

### A Genome-Wide Screen in Yeast Identifies New Regulators of ERK-Type MAPK Pathways

Strains used in the study are listed in Table 1. Three MAPK pathways in yeast require a subset of common components, including the Rho-type GTPase Cdc42, PAK Ste20, and MAPKKK Ste11, yet the pathways induce different responses [**Figure 1**, (Roberts et al. 2000; Maleri et al. 2004)]. A genetic screen was performed to identify regulators of ERK-type MAPK pathways in yeast. An ordered collection of overexpression plasmids (Gelperin et al. 2005) was examined for the induction of a MAPK pathway-dependent growth reporter [**Figure 2A**, *FUS1-HIS3*, (McCaffrey et al. 1987a; Horecka and Sprague 2000)]. The reporter provides a readout of two ERK-type MAPK pathways, mating and fMAPK (**Figure 1)**. However, cells lacking an intact mating pathway were evaluated (*ste4*Δ), which biases reporter activity towards the fMAPK pathway (Cullen et al. 2004). Specifically, high-throughput transformation was used to introduce 4416 plasmids into a wild-type yeast strain (**Figure 2A**) (Cullen et al. 2004) containing the *FUS1-HIS3* reporter. Colonies were transferred in 96-well format from S-D-URA media to S-GAL-URA media to induce overexpression of the genes. After 24h, cells were pinned from S-GAL-URA to S-GAL-URA (as a control for growth), S-GAL-URA-HIS, and S-GAL-URA-HIS containing 3-<u>a</u>mino-1, 2, 4-tri<u>a</u>zole (ATA) media (10 microliters of 2M in 25 ml plates). ATA is a competitive inhibitor of the His3 enzyme, and its inclusion allows for selection for high levels of reporter activity (Kanazawa et al. 1988). Genes that inhibited growth on S-GAL-URA-HIS (**Figure 2A**, Low Threshold) may when overexpressed dampen reporter activity and will be discussed elsewhere. Genes that induce growth on S-GAL-URA-HIS+ATA media may stimulate MAPK pathway activity due to transcriptional up-regulation of the growth reporter (**Figure 2A)**. These genes, in principle, have the potential to encode new MAPK pathway regulatory proteins.

**Table 1.**
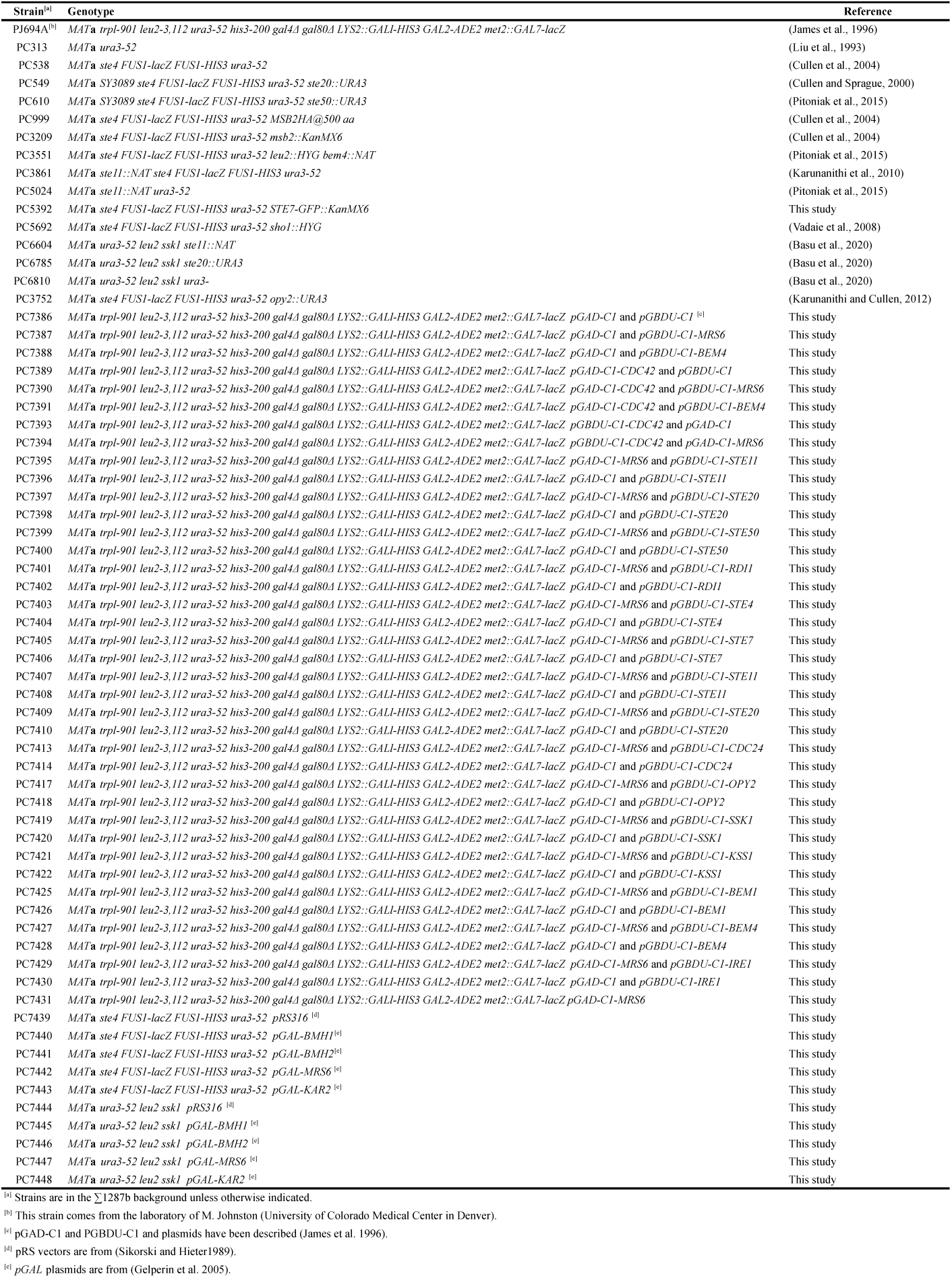
Yeast strains used in the study.

**Figure 1.**
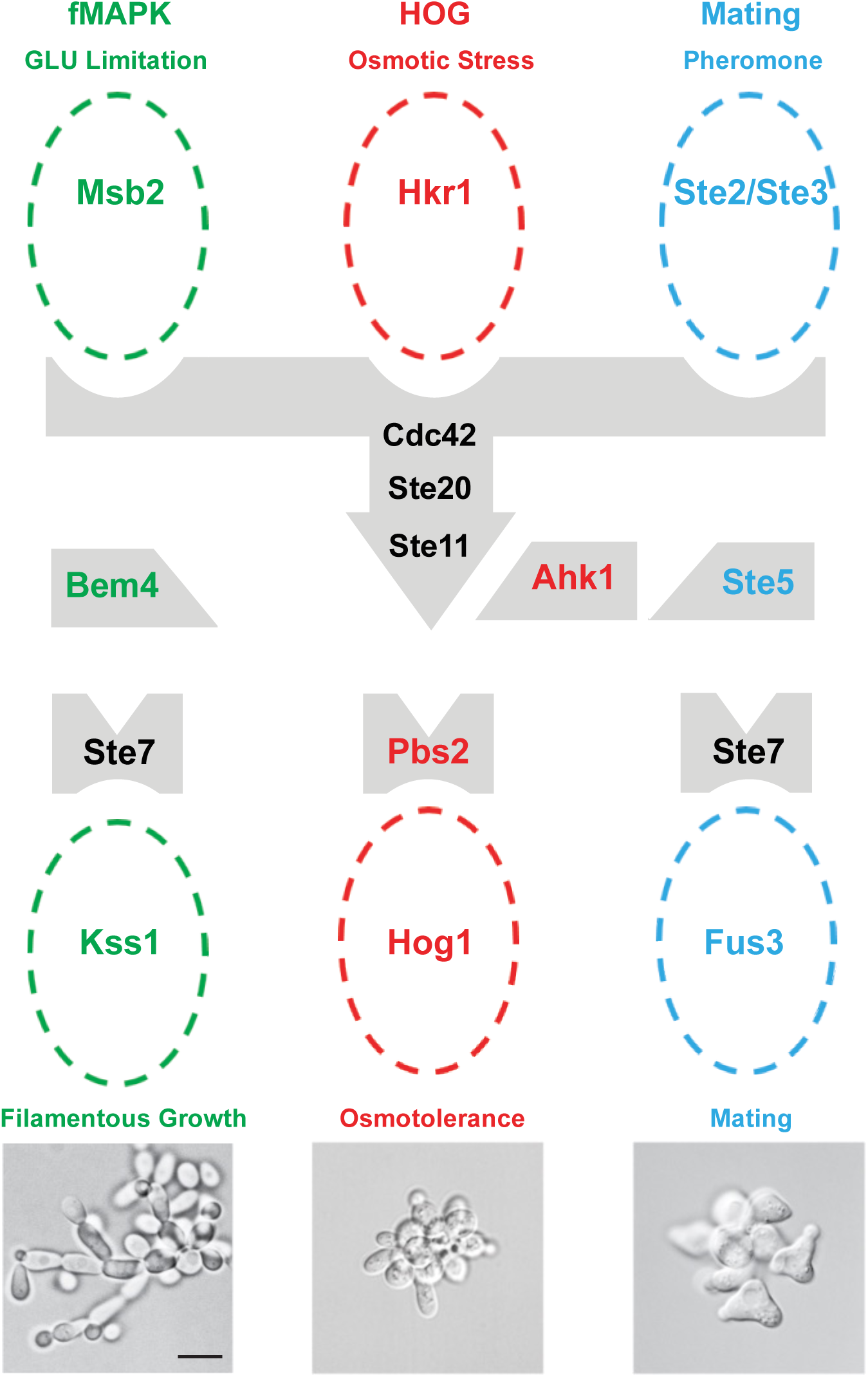
Three MAPK pathways in yeast share a subset of common components. Common components are shown in black, and pathway specific proteins are shown in color for the fMAPK (red), HOG (green), and mating (blue) pathways. Each pathway has a scaffold-type adaptor, Bem4 (Pitoniak et al. 2015), Pbs2 (Maeda et al. 1995b) and Ahk1 (Nishimura et al. 2016), and Ste5 (Chol et al. 1994), and MAP kinase. Cells undergo filamentous growth under nutrient-limiting conditions (left), cells do not change their morphology when exposed to YEP-GAL+1.0 M KCl salt (middle), YEP-GAL+1mg/ml α-factor stimulates an elongated cell shape or shmoo (right). Scale bar, 10 µm.

**Figure 2.**
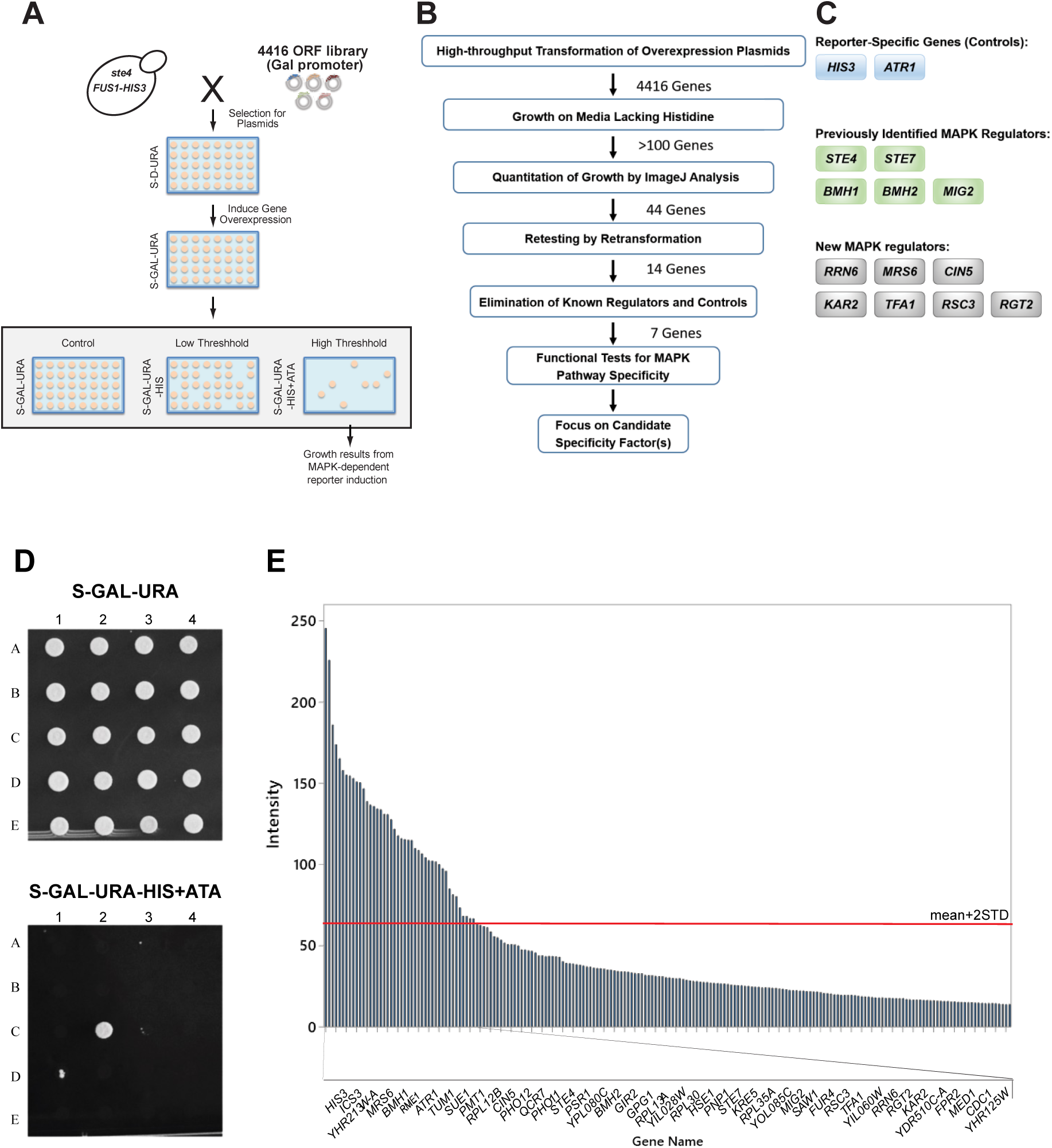
Genome-wide overexpression screen for new MAPK pathway regulatory proteins. **(A)** Diagram of the overexpression screen. An ordered collection of 4416 ORF overexpression plasmids covering ∼80% of yeast genome controlled by an inducible (*pGAL1*) promoter (circles (Gelperin et al. 2005)) were introduced into a *ste4 FUS1-HIS3* strain (PC999) by the high-throughput transformation. Transformants were generated by a microtiter plate method and pinned onto S-D-URA to select for plasmids. Overexpression of genes was accomplished by pinning colonies from S-D-URA to S-GAL-URA medium to induce overexpression of the genes. On the following day, cells were pinned to low threshold and high threshold (containing ATA, a competitive inhibitor of the His3 enzyme) media to identify genes that induce a MAPK pathway-dependent growth reporter (*FUS1-HIS3*) on media lacking histidine. The genes, which could overcome ATA, were identified as the candidates that, when overexpressed, can turn the pathway highly up (colored spots). **(B)** Pipeline for identifying functionally relevant MAPK pathway regulators. 44 genes were identified and prioritized for further analysis. The validation screen identified 14 robust genes from the initial screen. **(C)** The list of 14 genes that induced the MAPK pathway-dependent reporter, *FUS1-HIS3*, when overexpressed. Genes fell into three categories (see Table 2 for more details). **(D)** Example of a portion of one plate from the overexpression screen (the full screen is available in *Table S3*). The colony growing in the lower panel, C2, overexpresses *MRS6*. **(E)** The graph shows the results of the top genes identified by the overexpression. Colony growth on S-GAL-URA-HIS+ATA resulting from reporter (*FUS1-HIS3*) expression was measured by ImageJ analysis. Growth based on spot intensity and determined and plotted in the graph. The top 200 genes are shown. Forty-four genes passed a cut-off of mean+2STD (red bar) and are labelled here.

A scheme was employed to identify relevant MAPK pathway regulators, represented by a flowchart (**Figure 2B**). In the initial screen, >100 genes were identified that showed some growth on S-GAL-URA-HIS+ATA media (**Figure 2D**, *Table S3*). To quantitatively assess differences in growth, colony size was measured by ImageJ analysis [*Table S1*, (Schindelin et al. 2012)]. By applying a rigorous cut-off of the mean+2SD, 44 genes were identified that showed elevated MAPK reporter activity when overexpressed (**Figure 2, B** and **E**). To independently validate genes identified by the screen, plasmids containing candidate genes were re-transformed into wild-type cells and re-tested for reporter activity. Fourteen genes passed this validation step (**Figure 2, B** and **C**; see *Table S2* for the raw data).

**Table 2.**
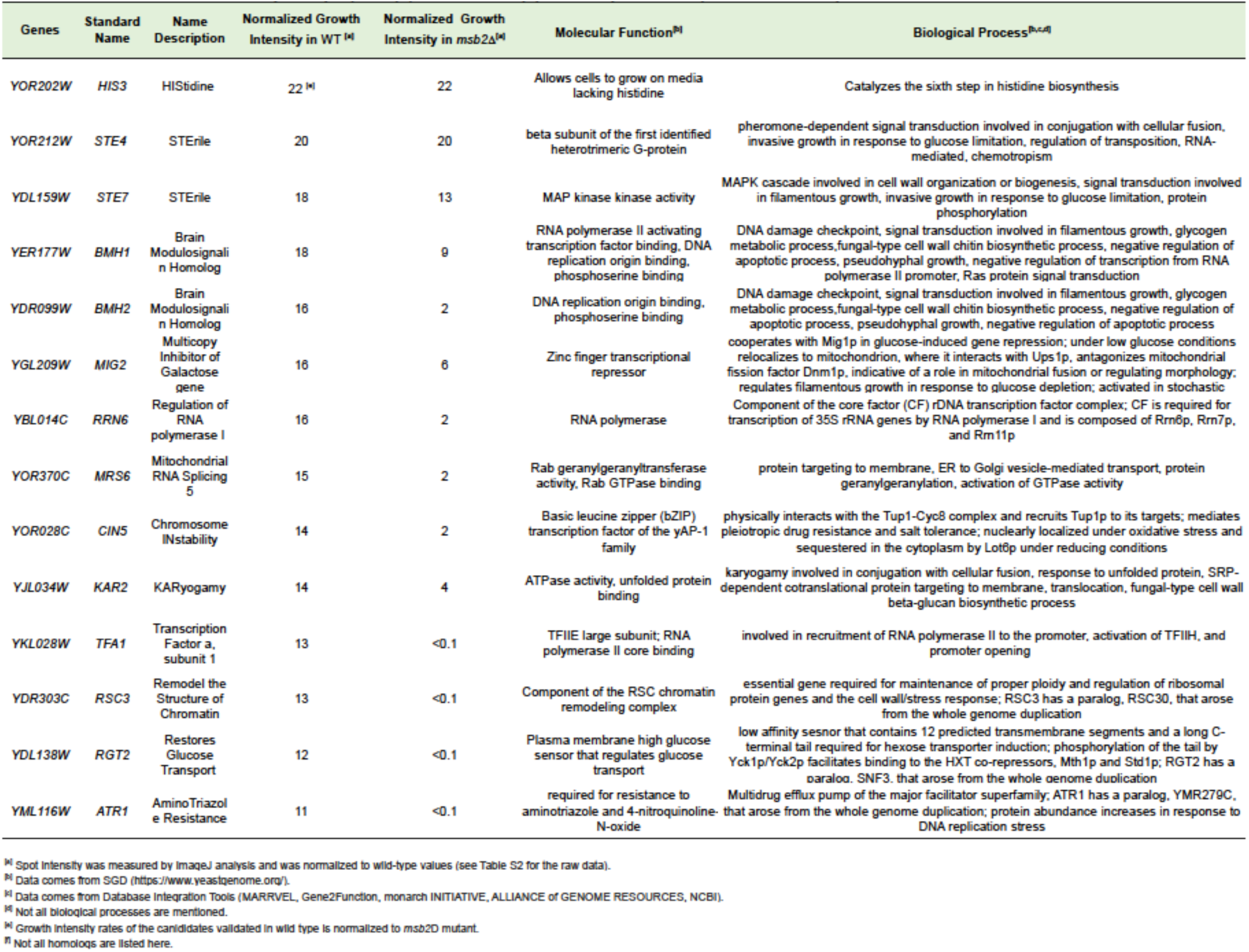
Functional classification of MAPK pathway regulatory quens identified by gene overexpression alongside human homologs.

The genes that passed the above criteria fell into three categories (**Figure 2C**, Table 2). The first category was metabolic controls. Two controls were identified, *HIS3*, which allows growth on media lacking histidine (Alifano et al. 1996), and *ATR1*, which encodes a multidrug efflux pump that confers ATA resistance (Kanazawa et al. 1988). The second category was known regulators of MAPK pathways. These included *STE4*, which regulates the mating pathway and complemented the signaling defect of the *ste4* mutant (Barr et al. 1996); *STE7*, the MAPKK that regulates the mating and fMAPK pathways (Neiman and Herskowitz 1994); *BMH1* and *BMH2*, which are members of the 14-3-3 family of proteins and are established regulators of the fMAPK pathway (Roberts et al. 1997), and *MIG2* a transcriptional repressor (Lutfiyya et al. 1998) that has been implicated in fMAPK pathway regulation (Karunanithi and Cullen 2012). Not all components of the fMAPK pathway were identified: *STE20, STE50*, and *STE11* were not present in the collection; *MSB2* and *CDC42* would not be expected to be identified as C-terminal fusions of the proteins, which occur in the library, are not functional in the fMAPK pathway; *OPY2* was identified but fell below the threshold for statistical significance, and *TEC1* does not induce the growth reporter. In its unphosphorylated form, the MAPK Kss1 would also not be expected to activate the reporter and may not be identified for this reason (Cook et al. 1997; Bardwell et al. 1998a; Bardwell et al. 1998b). *SHO1, BEM4*, and *STE12* were present in the collection but did not induce the reporter for reasons that have not been explored. The third category was potentially new MAPK pathway regulators. These included *RRN6, MRS6, CIN5, KAR2, TFA1, RSC3*, and *RGT2* (**Figure 2C**, *Table 2*).

To explore the characteristics of the genes identified by the screen, we used gene ontology (GO) annotations and database integration tools were used to identify the molecular and biological roles of proteins and determine whether they had mammalian homologs (Skrzypek and Hirschman 2011; Hu et al. 2017; Mungall et al. 2017; Wang et al. 2017; Coordinators 2018; 2020). Many of the identified genes had human homologs with established functions in diverse biological processes (Table 2). These included *BMH1, BMH2* (Roberts et al. 1997), *TFA1* (Feaver et al. 1994), *MRS6* (Alory and Balch 2000; Alory and Balch 2003), and *KAR2* (Rose et al. 1989). Moreover, the screen identified several essential genes (*MRS6, KAR2, TFA1*, and *RRN6*), and a set of paralogs (*BMH1* and *BMH2*), which might be missed in whole-genome deletion screens.

Many signaling pathways can influence the activity of the fMAPK pathway. One mechanism for this regulatory input comes from the regulation of the expression of the *MSB2* gene (Chavel et al. 2010). *MSB2* encodes the mucin-type glycoprotein that regulates the fMAPK pathway (Cullen et al. 2004). To determine whether these genes fall above or below Msb2 in their ability to stimulate MAPK pathway activity, candidates from the screen were examined for overexpression-dependent bypass the signaling defect of the *msb2*Δ mutant. A subset of the genes tested restored signaling in the *msb2*Δ mutant (Table 2, see *Table S2* for the raw data), which indicates that they function below the level of Msb2 in the MAPK pathway. We are interested in new regulators that, when overexpressed, bypass the signaling defect of the *msb2*Δ mutant (*MRS6, RRN6*, and *KAR2*), because of their potential to modulate MAPK pathway activity directly.

### Examining the Role of New MAPK Pathway Regulators in Polarity Reorganization during Filamentous Growth

One consequence of fMAPK pathway induction is enhanced polarized growth (Cullen et al. 2004). During filamentous growth, cells become highly polarized, which is evident by the elongated shapes of cells. The single-cell invasive growth assay (Cullen and Sprague 2000) was used to examine the polarized growth of a subset of candidate genes identified in the screen. Wild-type cells exist in the yeast form when grown in glucose (**Figure 3**, S-D-URA) and undergo filamentous growth when grown in the non-preferred carbon source galactose (**Figure 3**, S-GAL-URA). Cells lacking an intact fMAPK pathway are defective for filamentous growth by this assay (**Figure 3**, *ste20*Δ). Overexpression of *MRS6, BMH1, BMH2, KAR2*, and *TFA1* induced hyperpolarized growth. Specifically, the cells were longer and had irregular morphologies (**Figure 3**, arrows). Hyperpolarized morphologies result from a delay in the cell cycle, prolonging the period of polarized growth (Kron et al. 1994; Loeb et al. 1999; Madhani et al. 1999). This phenotype is consistent with the elevated activity of the fMAPK pathway. Indeed, this phenotype is distinct from activation of the HOG pathway, which shares components with the fMAPK pathway but does not induce a morphogenetic change when gets activated (Saito 2010a).

**Figure 3.**
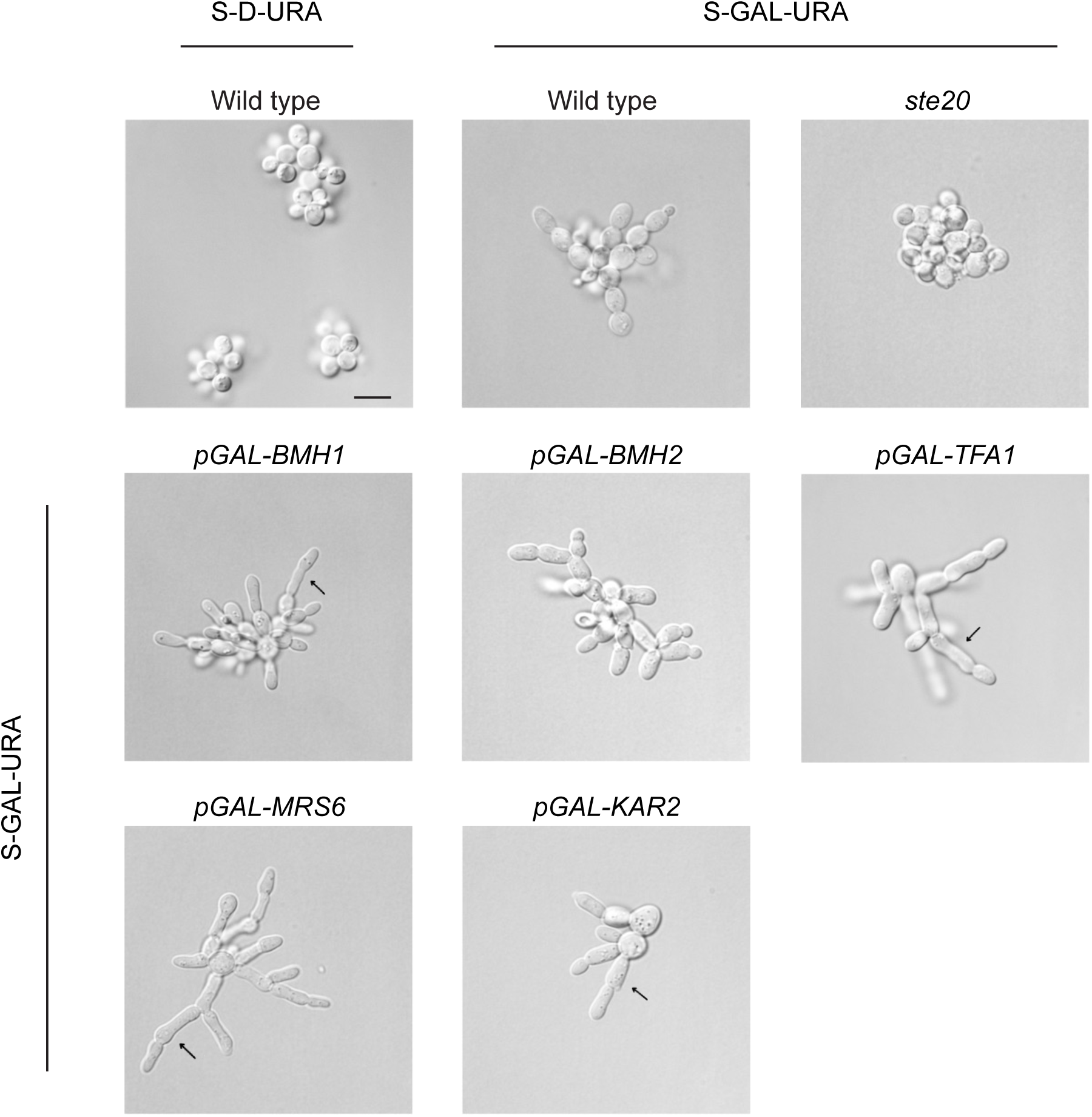
Morphological analysis of cells overexpressing genes that stimulate MAPK pathway signaling. Cell morphology of the indicated strains by the single-cell invasive growth assay by DIC microscopy at 100X magnification. Scale bar, 10 µm. As controls, wild-type cells were grown in glucose (Glu, S-D) and galactose (S-Gal) media, and the *ste20*Δ mutant was grown in S-Gal media. Overexpression of *MRS6, BMH1, BMH2, KAR2*, and *TFA1* induced hyperpolarized morphologies. Arrows show elongated cells making chains of filaments.

### Analysis of fMAPK Pathway Regulatory Proteins by Functional Tests for MAPK Pathways

Three MAPK pathways in yeast share components, including the Rho-type GTPase Cdc42, PAK Ste20, and MAPKKK Ste11 (**Figure 1**, (Roberts et al. 2000; Maleri et al. 2004)). As a complementary approach to assess the role of overexpression of *MRS6, BMH1, BMH2, KAR2*, and *TFA1* on these pathways, functional tests were performed that provide a readout of the three MAPK pathways. The plate-washing assay (PWA) measures the invasion of cells into the agar, which can be revealed by washing plates in a stream of water, and which is dependent on an intact fMAPK pathway (Roberts and Fink 1994b). Salt sensitivity was used to measure the activity of the HOG pathway (Maeda et al. 1994; Maeda et al. 1995b), and growth arrest by α-factor (halo assay) was used to measure the activity of the mating pathway (Sprague Jr et al. 1983). Growth on galactose (YEP-GAL) resulted in hyper-invasive growth for each of the candidate genes tested by the PWA (**Figure 4A**, green). Overexpression of *BMH1, BMH2*, and *MRS6* caused a growth defect on media containing salt (**Figure 4A**, red). This result was interesting because the fMAPK pathway functions antagonistically with the HOG pathway (Adhikari and Cullen 2014). Thus, it is plausible that elevated activation of the fMAPK pathway by overexpression of these genes might result in a dampened HOG response. Overexpression of these genes did not result in a defect in halo formation (**Figure 4A**, blue). Quantifying the data also supports the idea that some genes turn fMAPK pathway up, and turn HOG down (**Figure 4B)**. These results support the idea that *MRS6, BMH1, BMH2, KAR2*, and *TFA1* stimulate the activity of the fMAPK pathway and might potentially play a specific role in that pathway when these genes are overexpressed.

**Figure 4.**
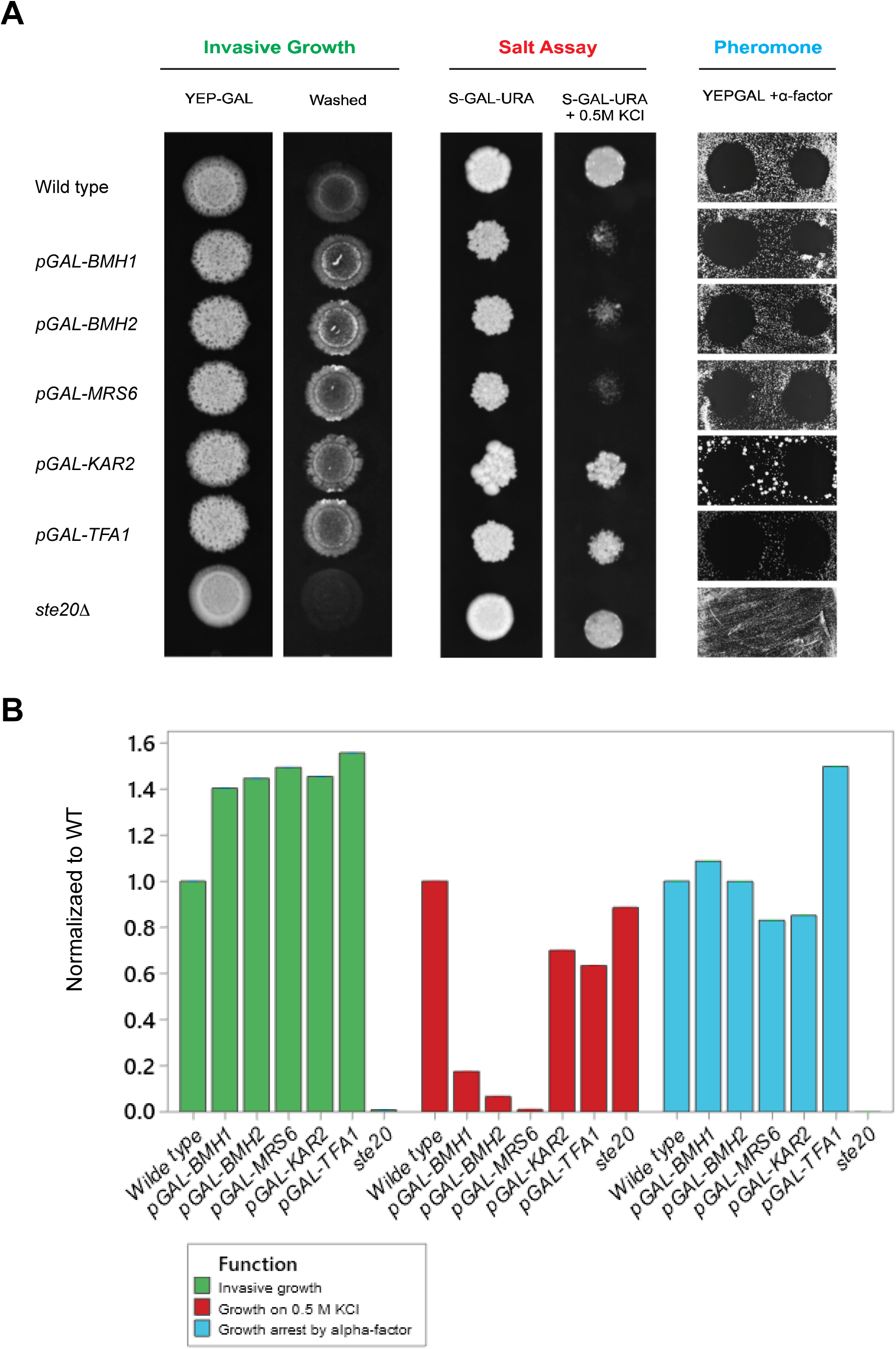
Phenotypic analysis of the role of overexpression of selected candidates on MAPK pathway activity. **(A)** Wild-type cells (PC6810) containing the indicated plasmids were grown in S-D-URA for 16 h and spotted onto the indicated media. For the PWA, cells were spotted onto YEP-GAL medium for 96 h. The plate was photographed (YEP-GAL), washed in a stream of water, and photographed again (Washed). To assess salt sensitivity, cells were spotted on S-GAL-URA and S-GAL-URA + 0.5 M KCl media for 72 h at 30°C. To determine sensitivity to α-factor, cells were spread onto S-GAL-URA plates. 10µl and 3µl drops of 1mg/ml α-factor were applied to the plates followed by incubation for 48 h. **(B)** plot shows the quantified data for the invasive growth, salt assay, and pheromone. Values normalized to wild type (WT) values, which were set to a value of 1.

### Mrs6 Overexpression Stimulates the fMAPK Pathway and Dampens the HOG Pathway

We focused on *MRS6* because it was one of the strongest hits from the screen (**Figure 2D**, the spot represents *MRS6*). Overexpression of *MRS6* also strongly induced polarized growth (**Figure 3**), induced hyper-invasive growth (**Figure 4**, green), and dampened the HOG pathway (**Figure 4**, red). Mrs6 is also an essential protein and, when overexpressed, bypassed the signaling defect of the *msb2* mutant (*Table 2*). Mrs6 is a Rab escort protein (Benito-Moreno et al. 1994) and has recently been identified as a modulator of the activity of the TOR pathway (Singh and Tyers 2009).

To explore the role of Mrs6 in regulating MAPK pathways, the phosphorylation of MAP kinases was examined, which provides a diagnostic readout of their activities. Using anti phospho p44-42 antibodies that detect the phosphorylated MAP kinases, Kss1 and Fus3, we found that overexpression of *MRS6* induced phosphorylation of Kss1 (**Figure 5A**, P∼Kss1). In comparison to wild type cells, where the levels of P∼Kss1 increased after 3 h growth in Gal and decreased after 7h, overexpression of *MRS6* caused a delay in the phosphorylation of Kss1, which was sustained until 11 h and then decreased (**Figure 5B**). This result indicates that *MRS6* alters the kinetics of the fMAPK pathway, in a manner that might be expected to stimulate fMAPK pathway activity.

**Figure 5.**
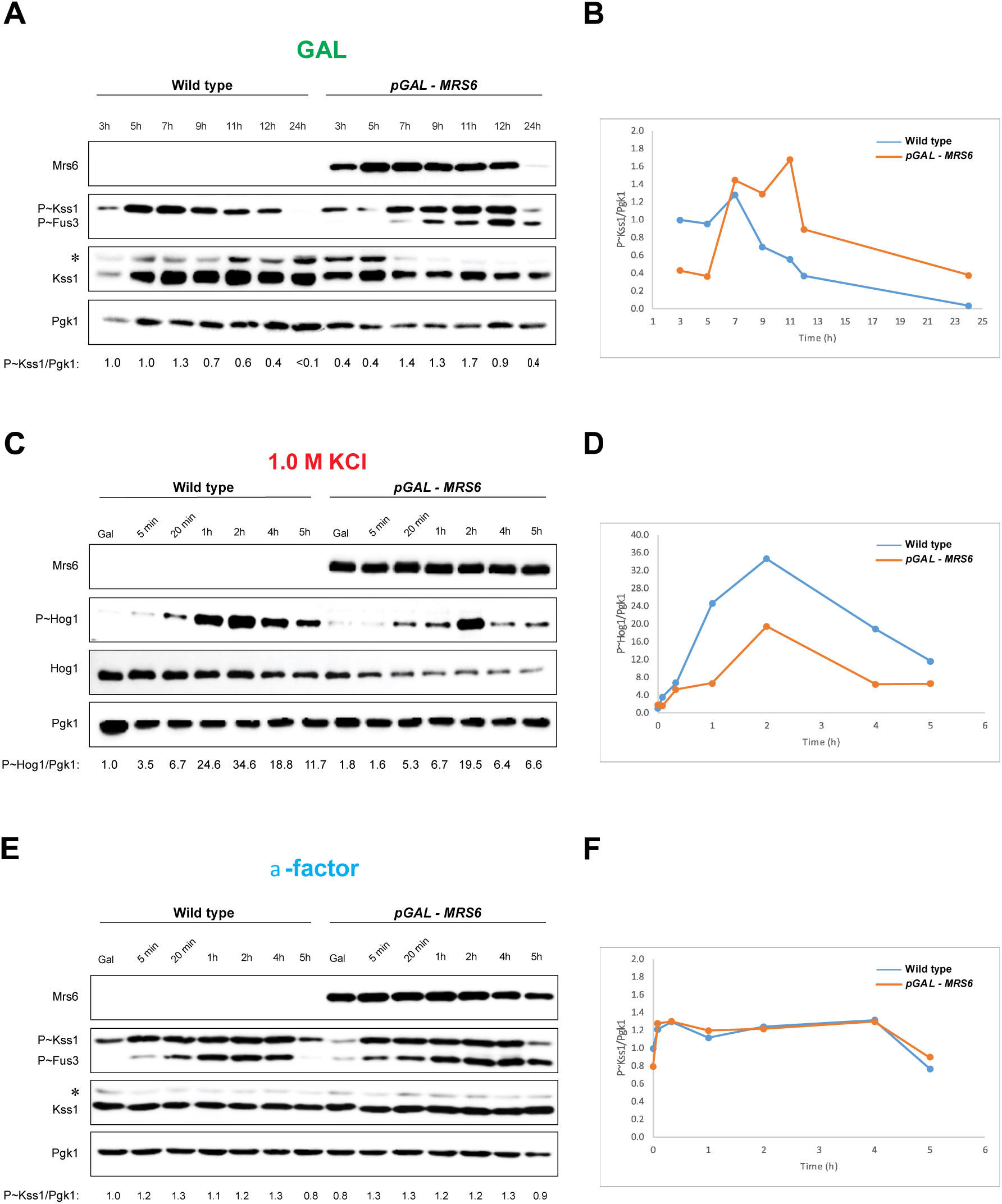
Impact of overexpression of Mrs6 on the fMAPK, HOG, and mating pathways. Wild-type cells (PC6810) and cells overexpressing *MRS6* (PC7447) were examined under conditions that induce MAPK pathway signaling. Cell extracts were evaluated by MAP kinase phosphorylation by immunoblot (IB) analysis. (**A**) Cells were grown in the non-preferred carbon source galactose (YEP-GAL), for the times indicated. Cell extracts were examined by IB analysis for P∼Kss1 and P∼Fus3 by p44/42 antibodies. Mrs6 proteins were detected at ∼91 kDa. Total Kss1 levels and Pgk1 (loading control, ∼45 kDa) also were assessed. The ratio of P∼Kss1 to Pgk1 normalized to wild-type values, which were set to a value of 1. (**B**) Graph visualizes the ratio of P∼Kss1 to Pgk1 for wild-type and *pGAL-MRS6*. **(C)** Cells were pre-grown in YEP-GAL for 4h following by growing in YEP-GAL medium containing 1.0 M KCl to examine P∼Hog1. (**D**) Graph showing P∼Hog1 to Pgk1 ratios, normalized to wild-type values, which were set to a value of 1. (**E**) Phosphorylation of Kss1 and Fus3 in response to pheromone. Cells were grown in YEP-GAL for 4h, and incubated in YEP-GAL medium containing 1mg/ml α-factor for the times indicated. Fus3 bands run in the same size as a degradation product of *MRS6*. (**F**) Graph showing P∼Kss1 to Pgk1, normalized to wild-type values, which were set to a value of 1.

By comparison, the terminal MAP kinase in the HOG cascade, Hog1, was not phosphorylated in response to salt in a manner that was influenced by *MRS6* overexpression (**Figure 5C**). In fact, overexpression of *MRS6* caused a modest reduction in HOG pathway activity (**Figure 5D**). These results match with the fact that overexpression of *MRS6* caused a growth defect on high-osmolarity media (**Figure 4**). Given that the fMAPK and HOG pathways can function antagonistically (Adhikari and Cullen 2014), our results suggest that *MRS6* may be a specific regulator of the fMAPK pathway. Overexpression of *MRS6* did not have a dramatic effect on the mating pathway (**Figure 5, E** and **F**). The main MAP kinase for the mating pathway, Fus3, is phosphorylated in response to pheromone. Although Fus3 phosphorylation was similar between wild-type cells and cells overexpressing *MRS6*, Fus3 migration overlapped with a degradation product of Mrs6 and was not used for quantitation. Therefore, *MRS6* overexpression led specifically to phosphorylation (activation) of the MAP kinase Kss1, which is consistent with a specific role for the protein in regulating the fMAPK pathway.

### Mrs6 Interacts with the Protein Kinases Ste20 and Ste7

To define how Mrs6 might specifically regulate the fMAPK pathway, genetic suppression analysis was performed. Genetic suppression analysis can allow the ordering of proteins into a pathway using gain- and loss-of-function alleles. p*GAL-MRS6* was introduced into mutants that lack fMAPK pathway components. Reporter induction by overexpression of Mrs6 was compared in cells lacking components of the fMAPK pathway (**Figure 1**). We looked at many components of fMAPK, including the *msb2*Δ, *sho1*Δ, *opy2*Δ, *ste20*Δ, *bem4*Δ, *ste50*Δ, and *ste11*Δ mutants. The results showed that Mrs6 overexpression partially bypassed the signaling defect of the *sho1*Δ mutant but not the *ste11*Δ mutant (*Fig. S1*, data shown for *sho1*Δ and *ste11*Δ). This experiment indicates that Mrs6 regulates the fMAPK pathway at or above the level of Ste11 in the fMAPK pathway. We also noticed that overexpression of *MRS6* induced a growth defect. The growth defect was separate from its induction of the fMAPK pathway, as it was seen in cells lacking fMAPK pathway components (*Fig. S2*).

To further define how Mrs6 regulates the fMAPK pathway, we analyzed the ability of Mrs6 to interact with fMAPK components by the two-hybrid system (Fields and Song 1989). Two-hybrid analysis can identify protein interactions *in vivo* by reconstitution of the binding and activation domains of fusion proteins to the Gal4 transcription factor, evaluated by a growth reporter (Fields and Song 1989). Two-hybrid analysis has proven to be a useful tool in detecting interactions in many biological systems, including the isolated domains of interacting proteins (Allen et al. 1995; Espenshade et al. 1995). The gene encoding Mrs6 was cloned into a two-hybrid vector (bait) and probed for interactions with a panel of proteins that regulate MAP kinase pathways. The analysis identified a robust interaction between Mrs6 and Ste20 (**Figure 6A**). Two-hybrid analysis also identified an interaction between Mrs6 and Ste7. Also, although we saw a very weak positive signal for the Ssk1 protein. Mrs6 did not associate with other components of fMAPK by two-hybrid analysis. Therefore, the two-hybrid analysis may provide an explanation for how *MRS6* promotes fMAPK signaling, which includes the kinases <u>Ste20</u>, Ste11, <u>Ste7</u>, and Kss1, but not the HOG pathway, which includes the kinases <u>Ste20</u>, Ste11, Pbs2, and Hog1 (**Figure 6B**).

**Figure 6.**
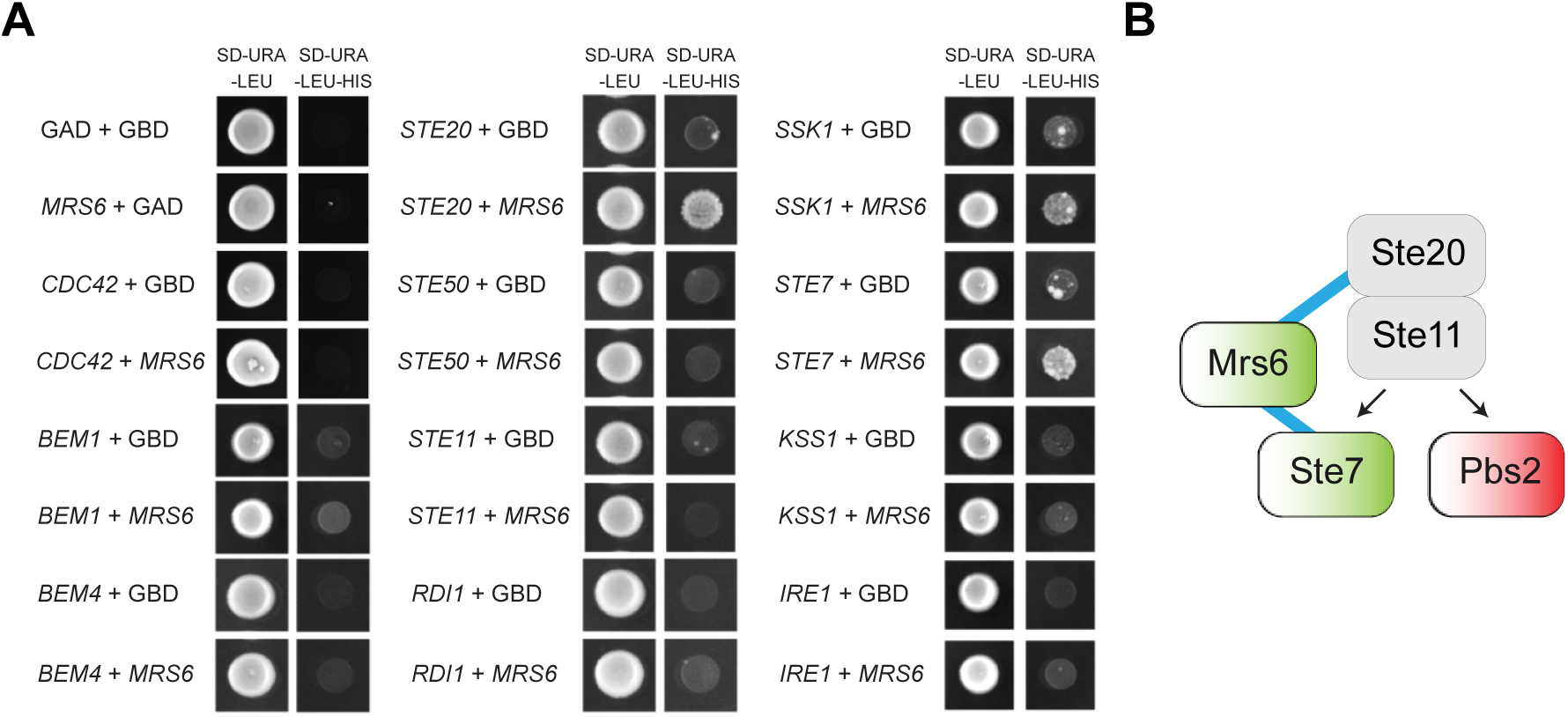
Two-hybrid analysis between Mrs6 and proteins that regulate fMAPK pathway. In the panels, GAD refers to pGAD-C1, and GBD refers to pGBDU-C1. (**A**) Cells were grown on S-D-URA-LEU to maintain selection for the bait and prey plasmids. Growth on medium lacking histidine (S-D-URA-LEU-HIS) displayed an interaction of Mrs6 with MAPKKK kinase Ste20, and an interaction between Ste7 and Mrs6. Based on two-hybrid analysis, Mrs6 did not associate with other components of fMAPK. (**B**) The two-hybrid analysis may provide an explanation for how *MRS6* promotes fMAPK signalling through the kinases <u>Ste20</u>, Ste11, <u>Ste7</u>, and Kss1, but it is not regulating the HOG pathway, which requires the kinases <u>Ste20</u>, Ste11, Pbs2, and Hog1.

## DISCUSSION

MAPK pathways regulate diverse cellular responses and are controlled by an expanding repertoire of regulatory proteins. In this study, we uncovered new regulators of an ERK-type MAPK pathway in yeast. We screened a *S. cerevisiae* library of covering 80% of the genome for genes that, when overexpressed, induce a MAPK-dependent growth reporter. We identified 12 regulatory genes of the MAPK pathway and two metabolic controls. The seven new genes identified in this study as MAPK pathway regulators provide a platform for exploring how a different cellular processes connect to and regulate MAPK pathways. We followed up on one of these candidates and showed by a combination of genetic and biochemical approaches that Mrs6 regulates the MAPK pathway that controls filamentous growth. Furthermore, Mrs6 interacts with kinases that regulate that pathway and might play a role in pathway specificity.

*MRS6* is an essential gene that may regulate the fMAPK pathway in several ways. One way might be through its role in regulating protein trafficking. Rab-type GTPases regulate protein trafficking (Bauer et al. 1996; Miaczynska et al. 1997; Lamber et al. 2019). Mrs6 is a Rab escort protein (REP) that makes a complex with the Rab GTPases Ypt1, Sec4, Ypt6, Vps21 (Fujimura et al. 1994; Bialek□Wyrzykowska et al. 2000; Sidorovitch et al. 2002). Given that Cdc42p is itself a component of the exocyst complex (TerBush et al. 1996; Zhang et al. 2001), it is possible that Mrs6 might contribute to the delivery of Cdc42 or other fMAPK pathway components to the plasma membrane. Similarly, Mrs6 may regulate the assembly of the fMAPK signaling complex and/or its function in the secretory pathway. Ypt1 also regulates the UPR by promoting the decay of HAC1 RNA (Tsvetanova et al. 2012). Interestingly, Msb2 and the fMAPK pathway are regulated by the UPR (Adhikari et al. 2015). Perhaps some of the regulators identified in this study connect the MAPK pathway to the UPR pathway. In a related study, we found that *BMH1* and *BMH2* showed a connection to the UPR but not *MRS6* (Jamalzadeh et. al, unpublished data).

Mrs6 facilitates geranylgeranylation for the prenylation of Ypt1 at the Golgi through the Bet2 and Bet3 enzymes (Witter and Poulter 1996). In the fMAPK pathway, the Rho-type GTPase Cdc42 is modified by lipid geranyl groups (by Cdc43); thus, Mrs6 may impact the lipid modification of Cdc42. However, Mrs6 did not associate with Cdc42 by two-hybrid analysis. Rather, a two-hybrid analysis showed that Mrs6 interacts with Ste20 and Ste7. Ste20 is the PAK kinase that regulates the fMAPK pathway (Roberts and Fink 1994b; Peter et al. 1996; Leberer et al. 1997). It has been reported that Ste20 is recruited by a complex containing Cdc42 and Cdc24 to the membrane (Pesce et al. 2016). Taken together, Mrs6 may regulate Cdc42 through the recruitment of Ste20.

Mrs6 interacts with Ste7 (a MAPKK). Given that *MRS6* specifically stimulates the fMAPK pathway, we speculate that Mrs6 might function as a scaffold to facilitate interactions among members of the kinase cascade (**Figure 6B)**. In support of this possibility, overexpression of *MRS6* dampened the activity of the HOG pathway. Therefore, Mrs6 may help minimize cross-talk with other MAP kinase cascades and thus ensure the integrity of the filamentous growth response.

In a separate study, *MRS6* was shown to regulate the TORC1 pathway through *SFP1* to control ribosome biogenesis (Lempiainen et al. 2009; Singh and Tyers 2009). TOR is a master regulatory pathway of cell growth and nutrient sensing (Kunkel et al. 2019). TOR’s activator, GOLPH3, has been identified recently as an oncogene in many human cancers (Scott et al. 2009). Hence, the identification of Mrs6 as a key regulator of the fMAPK pathway in yeast raises the possibility that REP1/REP2 may link fMAPK signaling to the TOR pathway and to the secretory system in higher organisms.

In mammalian cells, *MRS6* homolog encoded by CHM, which is the human Rab escort proteins REP1/CHM or REP2/CHML and share 50% sequence identity with Mrs6 (Alory and Balch 2003). CHM is a disease of the retina, which causes progressive vision loss (Alory and Balch 2001). Furthermore, REP1/CHM has been shown to regulate the epidermal growth factor receptor (EGFR) through the transcription factor STAT3. EGRF is also a major regulator of the Grb-SOS-RAS-MEK-ERK pathway, which is commonly misregulated in cancer cells (Yuan et al. 2004). Given that EGFR also signals through RAS-MEK-ERK (Zebisch et al. 2007; Li et al. 2016), our screen may have identified a new and general regulator of ERK-type MAPK pathways.

## MATERIALS AND METHODS

### Strains and Plasmids

Strains used in the study are listed in Table 1. Strains were cultured in yeast extract and peptone (YEP) media (1% yeast extract and 2% bactopeptone) with a source of carbon [2% glucose (D) or 2% galactose (GAL)] for growth in liquid culture or 2% agar for growth in semi-solid agar media. All experiments were carried out at 30°C unless otherwise specified. Synthetic complete (S) medium was used for maintaining selection for plasmids. Bacterial cultures of *Escherichia coli* were proliferated in LB+CARB media (carbenicillin) by standard methods (Sambrook et al. 1989). The pRS plasmids (pRS315 and pRS316) have been described (Sikorski and Hieter 1989). To construct two-hybrid plasmids, plasmids pGAD-C1 and pGBDU-C1 were used (James et al. 1996).

### Analysis of a Gene Overexpression Collection for Altered Activity of a MAPK Pathway-Dependent Growth Reporter

A microtiter-based high throughput transformation method (Gietz and Schiestl 2007) was used to introduce a collection of ∼4,500 plasmids, each overexpressing a different yeast gene (Gelperin et al. 2005) into strain (PC999). Transformants were screened for Msb2-HA secretion as described (Chavel et al. 2010) and the activity of the fMAPK pathway in this study. Specifically, transformants were pinned onto S-D-URA to select for plasmids. Colonies that grew onto S-D-URA were then pinned to S-GAL-URA to induce gene overexpression. From S-GAL-URA, cells were pinned onto S-GAL-URA, S-GAL-URA-HIS, and S-GAL-URA-HIS+ATA to identify positive regulators of the fMAPK pathway. Colonies that grew on S-GAL-URA-HIS+ATA media resulted from elevated fMAPK pathway activity due to the up-regulation of the growth reporter (*FUS1-HIS3*).

### Genome-Wide Screen and Data Analysis

The growth of 4416 genes was examined from 46 plates (raw data is available in Table S3). Not all of the genes from the collection were analyzed. This could be resulted because of the failure of some plasmids to be transformed and contamination on several plates. ImageJ analysis (https://imagej.nih.gov/ij/) was used to quantify colony growth. Images of the plates from the screen were converted to 8-bit and inverted. A threshold adjustment was performed, followed by analysis by the DNA microarray plugin to measure spot intensity for each colony (*Table S1*). Outputs from ImageJ were saved as cvs format for additional analysis.

A MATLAB script was written to identify growth that was statistically significant. A cut-off of mean+2STD identified the top 3% of genes that, when overexpressed, showed growth that was above background. Validation of candidates was performed by re-transformation of plasmids containing genes, by standard transformations (Gietz et al. 1995), into a wild-type strain (PC6021) and testing for reporter induction by growth on S-GAL-URA-HIS+ATA media (*Table S2*). The same plasmids were also transformed into the *msb2*Δ mutant (PC3209) to determine the bypass of that regulator of the pathway (*Table S2*). Database Integration Tools were used for further describing the identified genes’ characteristics and their orthologs in a concise manner (Skrzypek and Hirschman 2011; Hu et al. 2017; Mungall et al. 2017; Wang et al. 2017; Coordinators 2018; 2020).

### Microscopy

Differential interference contrast (DIC) microscopy was performed at 100X using an Axioplan 2 fluorescent microscope (Zeiss) with a Plan-Apochromat 100X/1.4 (oil) objective (N.A. 1.4) (coverslip 0.17) (Zeiss). Digital images were obtained with the Axiocam MRm camera (Zeiss) and Axiovision 4.4 software (Zeiss). Adobe Photoshop was used for brightness and contrast adjustments. Polarized cells were assigned by examining cells over multiple focal planes by DIC.

### Functional Assays for MAPK Pathway Activity

Cell morphology was assessed by the single-cell invasive growth assay (Cullen and Sprague 2000). Invasive growth was assessed by the PWA (Roberts and Fink 1994b). For the PWA, equal concentrations of cells were spotted onto YEP-GAL media. The activity of the HOG pathway was assessed by growth on high-osmolarity media. Equal concentrations of cells were spotted onto S-GAL-URA and S-GAL-URA+0.5M KCl media. Halo assays were performed as described (Jenness et al. 1987). Cells were spotted onto YEP-GAL media followed by spotting 3 µl and 10 µl α-factor (1mg/ml) on the plate. Plates were incubated at 30°C and photographed at 24 h and 48 h. The single-cell invasive growth assay was performed as described (Cullen and Sprague 2000).

### Phospho-Immunoblot Analysis

Phosphorylation of different MAP kinases in response to different stimuli was examined as described (Lee and Dohlman 2008; Basu et al. 2016). Cells were grown to mid-log phase from a saturated culture in YEP-D or YEP-GAL media for 4 h. Cells were washed and sub-cultured into YEP-GAL, YEP-GAL with 1.0 M KCl, or YEP-GAL with α-factor (1mg/ml). Cells were collected at various times by centrifugation, washed once, and stored at −80 °C. Proteins were extracted by trichloroacetic acid precipitation (TCA) and resuspended in 0.15 ml sample buffer by heating to 90°C. Protein samples were separated by sodium dodecyl sulfate-polyacrylamide gel electrophoresis (SDS-PAGE) (10% acrylamide). Proteins were transferred from polyacrylamide to nitrocellulose membranes (AmershamTM ProtranTM Premium 0.45 µm NC, GE Healthcare Life sciences, 10600003) by electrotransfer (Bio-Rad laboratories Inc.). Membranes were blocked with 5% BSA in 1X TBST (10 mM TRIS-HCl pH 8, 150 mM NaCl, 0.05% Tween 20).

Phosphorylation of mating and fMAPK pathways (P∼Kss1 and P∼Fus3) was investigated with p44-42 antibody (Cell Signaling Technology, Danvers, MA, 4370) at a dilution of 1:10000 to detect ERK-type MAP kinases. Phosphorylated Hog1 was detected using a 1:10000 dilution of α-phospho p38 antibody (Santa Cruz Biotechnology, Santa Cruz CA; #yC-20). Total Kss1 was detected with α-Kss1 antibodies (Santa Cruz Biotechnology, Santa Cruz, CA; #6775) at a 1:5,000 dilution. Total Hog1 was detected with α-Hog1 antibodies at a 1:5,000 dilution. Membranes were incubated 16 h with primary antibodies in 1X TBST with 5% BSA at 4°C. Control membranes were incubated 16 h in Pgk1 antibodies in 1X TBST with 5% non-fat dried milk at 4°C. To detect the primary antibodies, secondary antibodies of Mouse α-Pgk1 at a 1:5,000 dilution (Novex, 459250), goat anti-rabbit IgG-HRP at a 1:10,000 dilution (Jackson ImmunoResearch Laboratories, Inc., West Grove, PA, 111-035-144), and Goat α-mouse secondary (Bio-Rad Laboratories, Hercules, CA, 1706516) at a 1:5,000 dilution were used within milk as blocking buffer. Blots were visualized by chemiluminescence using a Bio-Rad ChemiDoc XRS+ system (Bio-Rad, 1708265). Image Lab Software (Bio-Rad, Inc.) was applied to analyze the band intensity.

### Genetic Suppression Analysis

Control (pRS316) and *pGAL-MRS6* plasmids were transformed into wild-type strain (PC538) and MAPK pathway mutants. These included *msb2*Δ (PC3209), *sho1*Δ (PC5692), *opy2*Δ (PC3752), *ste20*Δ (PC5692), *bem4*Δ (PC3551), *ste50*Δ (PC610), and *ste11*Δ (PC3861) mutants. Cells were grown on S-GAL-URA and S-GAL-URA-HIS to evaluate growth, which we infer to represent bypass of the mutant phenotype.

### Cloning the MRS6 Gene into Two-Hybrid Plasmids

The *MRS6* gene was cloned into the pGAD-C1 and pGBDU-C1 vectors in the following way. The *MRS6* gene was amplified by PCR using the forward primer 5’-ATGCATCGATATGTTAAGTCCTGAACGTAGACC-3’ and reverse primer of 5’-ATGCGTCGACTCATATCTCCATTTCACCTACAAATTC-3’. The PCR product was purified with QIAquick PCR Purification Kit, Qiagen, CA#28106. The PCR product and pGAD-C1 vector were digested with ClaI (5’-ATCGAT-3’, New England BioLabs Inc., MA, CA#R0197S) and SalI (5’-GTCGAC-3’, New England Biolabs Inc., MA, CA#R3138S) restriction enzymes. Digested insert and vector DNAs were run on a 1% agarose gel containing ethidium bromide. Bands were extracted from the gel using the QIAquick Gel Extraction Kit, Qiagen (CA#28704). A quick Ligation Kit (New England Biolabs Inc., MA, CA#M200l) was used for ligating the insert and vector. The ligation mixture was transformed into *E. coli* (One-Shot MAX Efficiency DH5α-T1 Competent Cells, ThermoFisher, CA# 12297016), followed by plating on LB+Carb plates. The plates were incubated at 37°C for 24 h. Transformants were confirmed by digestion with ClaI, and SalI Plasmids were sequenced at the Roswell Park Sequencing facility (Roswell Park Cancer Institute, Buffalo, NY).

### Two-Hybrid Assay

Two-hybrid constructs (pGBDU-C1 bait and pGAD-C1 prey) and empty vectors as controls were introduced into strain PJ694A (PC284) (James et al. 1996) using the lithium acetate transformation standard protocols (Gietz and Woods 2002). Transformants were selected on S-D media lacking uracil (URA) and leucine (LEU) to maintain selection for plasmids. Protein-protein interactions were screened by spotting cells onto S-D-URA-LEU media that was also lacking histidine (HIS) and containing ATA. Growth in this media results from the induction of a two-hybrid transcriptional reporter as the readout of protein-protein interactions.

## ABBREVIATIONS

ATA: 3-Amino-1, 2, 4-triazole
CARB: carbenicillin
CHM: choroideremia
CHML: choroideremia-like
D: dextrose
DIC: differential interference contrast
*E. coli*: *Escherichia coli*
ERK: extracellular-signal-regulated kinase
GAL: galactose
GLU: glucose
GAD: Gal4 activation domain
GBD: Gal4 binding domain
GDI: GDP-dissociation inhibitor
GDP: guanine nucleotide diphosphate
GGPP: geranylgeranyl pyrophosphate
GO: gene ontology
GTP: guanine nucleotide triphosphate
HIS: histidine
HOG: high osmolarity glycerol response
LEU: leucine
MAPK: mitogen activated protein kinase
MAPKKK: mitogen activated protein kinase kinase kinase
MEKK: MAP kinase kinase kinase
MEK: MAP kinase kinase
PAK: p21-activated protein kinase
PWA: plate washing assay
RabGDI: RabGDPdissociation inhibitor
RabGGTase: Rab geranylgeranyl transferase
RCR: REP-conserved region
REP: Rab escort protein
SCR: structurally conserved region
Rho: Ras homology
SDS-PAGE: sodium dodecyl sulfate-polyacrylamide gel electrophoresis
STD: standard deviation
S: synthetic
TCA: trichloroacetic acid
URA: uracil
WT: wild type
YEP: yeast extract and peptone.

## ACKNOWLEDGMENTS

Thanks to Heather Dionne for performing the screen. Thanks to previous and current members of the Cullen laboratory, especially Dr. Beatriz Gonzalez and Aditi Prabhakar, for technical assistance and insightful discussions through the course of this work. This work was supported by a grant from the NIH (GM098629).

The authors have no conflicts of interest to disclose.

The authors have no competing interests in the study.

